# UBQLN2 restrains the domesticated retrotransposon PEG10 to maintain neuronal health in ALS

**DOI:** 10.1101/2022.03.25.485837

**Authors:** Holly H. Black, Julia E. Roberts, Shannon N. Leslie, Will Campodonico, Christopher C. Ebmeier, Cristina I. Lau, Alexandra M. Whiteley

## Abstract

Amyotrophic Lateral Sclerosis (ALS) is a fatal, neurodegenerative disease characterized by progressive motor neuron dysfunction and loss. A portion of ALS is caused by mutation of the proteasome shuttle factor *Ubiquilin 2* (*UBQLN2)*, but the molecular pathway leading from UBQLN2 dysfunction to disease remains unclear. Here, we demonstrate a function of UBQLN2 in regulating activity of the domesticated gag-pol retrotransposon ‘paternally expressed gene 10’ (PEG10) in human cells and tissues. In cells, the PEG10 gag-pol protein cleaves itself in a mechanism reminiscent of retrotransposon self-processing to generate a liberated ‘nucleocapsid’ fragment, which uniquely localizes to the nucleus and changes expression of genes involved in axon remodeling. In spinal cord tissue from ALS patients, PEG10 gag-pol is elevated compared to healthy controls. These findings implicate the retrotransposon-like activity of PEG10 as a contributing mechanism in ALS through regulation of gene expression, and restraint of PEG10 as a primary function of UBQLN2.

## Introduction

Amyotrophic Lateral Sclerosis (ALS) is a fatal neurodegenerative disease which typically presents in mid-life and is characterized by a progressive loss of motor function (Brown and Al-Chalabi, 2017). Of all ALS cases, 90% are sporadic (sALS), while the remaining 10% are familial (fALS) and can be traced to mutations in a variety of genes, including the proteasome shuttle factor *Ubiquilin 2* (*UBQLN2)* (Deng et al., 2011; Gorrie et al., 2014; Williams et al., 2012). Animal models of fALS have led to discoveries that are broadly applicable to both fALS and sALS, including the involvement of oxidative stress, RNA binding proteins, and protein aggregation in ALS-mediated neuronal dysfunction (Ferraiuolo et al., 2011; Peters et al., 2015); however, the specific molecular pathways that lead to disease remain poorly understood.

In a previous global proteomic study, animal models of *UBQLN2*-mediated fALS revealed a dramatic accumulation of the domesticated retrotransposon ‘Paternally Expressed Gene 10’ (PEG10) in diseased tissue (Whiteley et al., 2021). Domesticated retrotransposons encode virus-like proteins that have lost pathological effects and have evolved adaptive functions (Volff, 2006). PEG10, which is necessary for placental development (Ono et al., 2006), is one of a family of domesticated retrotransposon genes derived from the Sushi-ichi lineage of Ty3/Gypsy LTR retrotransposons and codes for both *gag* and *pol* domains separated by a programmed ribosome frameshifting site (Brandt et al., 2005). Use of the ribosomal frameshift occurs with high efficiency, resulting in two cellular pools of PEG10 protein: gag, and gag-pol (Clark et al., 2007; Lux et al., 2010; Manktelow, 2005). The overwhelming share of research on PEG10 has focused on contributions to placental development (Abed et al., 2019; Ono et al., 2006) and cancer progression (Akamatsu et al., 2015; Kim et al., 2019); here, we describe a novel role in neurodegenerative disease.

## Results

### UBQLN2 exclusively regulates the frameshifted gag-pol PEG10

*UBQLN2* is one of five human Ubiquilin (*UBQLN*) genes which facilitate proteasomal degradation of ‘client’ proteins (Hjerpe et al., 2016; Itakura et al., 2016; Lee and Brown, 2012; Suzuki and Kawahara, 2016; Whiteley et al., 2017; Zheng et al., 2020) via an N-terminal protein domain which binds to the proteasome (Finley, 2009; Saeki, 2017), and a C-terminal domain which binds to ubiquitin (Zhang et al., 2008). All five *UBQLNs* have similar protein domain architecture and amino acid sequences, leading to the hypothesis that UBQLNs may have shared client populations. However, *UBQLN2* is unique for its enriched expression in neural tissues (Marín, 2014) and for containing a small, proline-rich PXX repeat region that is commonly mutated in *UBQLN2*-mediated fALS (Deng et al., 2011). To test the specificity of the relationship between UBQLN2 and PEG10, human embryonic stem cells (hESCs) lacking *UBQLN1*, *UBQLN2*, or *UBQLN4* genes (Figure S1A) were probed by western blot for endogenous PEG10 protein expression (Figure 1A). Only *UBQLN2*^-/-^ hESCs demonstrated an increase in PEG10 protein; furthermore, only the gag-pol form of PEG10 accumulated, while the gag form remained unchanged (Figure 1B-C).

**Figure 1:**
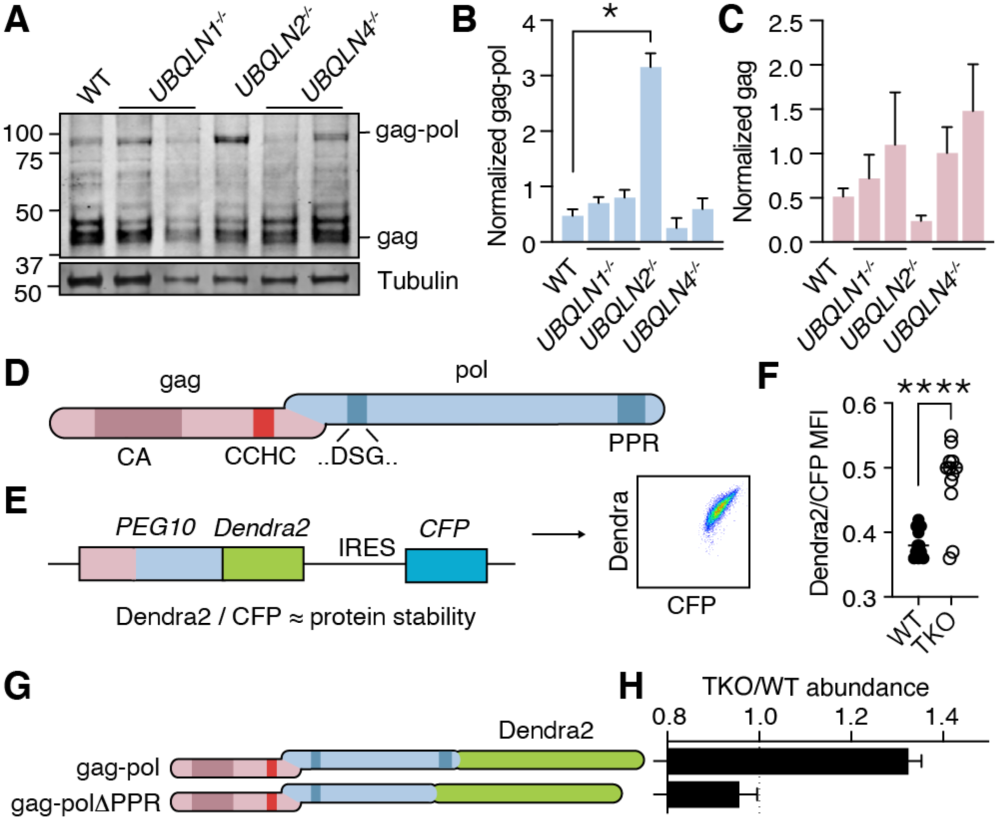
UBQLN2 regulates PEG10 gag-pol abundance. **(A)** Human ESCs had individual *UBQLN* genes deleted by CRISPR gene editing and clones were probed by western blot for endogenous PEG10 protein. Full-length gag-pol protein accumulates only upon *UBQLN2* loss. n=3 independent experiments. **(B-C)** Quantification of gag-pol (B) and gag (C) abundance in hESC cell lines of (A). PEG10 protein was normalized to Tubulin, then normalized to the average intensity for each individual experiment. n = 3 independent experiments, and significance was determined by multiple comparisons test. No differences in gag (C) were detected with an ordinary one-way ANOVA. **(D)** Schematic of PEG10 protein. The first reading frame (gag) contains a capsid-like (CA) region, as well as a retroviral zinc finger (‘CCHC’). The pol-like sequence contains a retroviral-type aspartic protease with one active site ‘DSG’ motif, as well as a C-terminal polyproline repeat domain (PPR). **(E)** Schematic of PEG10 protein abundance reporter. PEG10 is fused at the 3’ end to Dendra2, followed by an IRES-CFP. Right: example dot plot showing Dendra2 and CFP signal in transfected cells. **(F)** Dendra2 over CFP MFI ratio for PEG10 in WT and *UBQLN1*, *2*, and *4* ‘TKO’ HEK293 cells. Shown is mean ± SEM of MFI ratio from four independent experiments with triplicate transfection wells. Unpaired comparison between WT and TKO Dendra2/CFP ratios was performed by Student’s t-test. **(G)** WT and truncation mutant of PEG10 gag-pol fused to the fluorophore Dendra2. ΔPPR is missing the last 27 amino acids containing polyproline repeat. **(H)** Protein abundance of PEG10-Dendra2 fusions was determined for WT and TKO cells by flow cytometry. Values over 1.0 indicate dependence on *UBQLNs* for restriction. n = 5 independent experiments.

As UBQLN2 selectively regulated the gag-pol form of PEG10, we hypothesized that a unique region of the pol domain rendered it dependent on UBQLN2 for its degradation. The gag region contains a retroviral capsid domain and CCHC-type zinc finger (Figure 1D). The pol region of PEG10 is less well understood, but contains an aspartic protease domain (Clark et al., 2007) and a 27 AA C-terminal polyproline repeat (PPR) region containing twelve prolines in tandem, and 18 in total (Clark et al., 2007) (Figure 1D). To identify the region of PEG10 that defines its reliance on UBQLN2, either PEG10 gag-pol, or a construct lacking the C-terminal PPR, was fused to the fluorescent protein Dendra2 (Klementieva et al., 2016), followed by an IRES-CFP cassette, to generate a transfection-controlled measure of protein abundance (Figure 1E, Figure S1B). PEG10-reporter constructs were then transfected into WT and *UBQLN1*, *2*, and *4* triple knockout (‘TKO’) HEK293 cells (Itakura et al., 2016) to examine the abundance of PEG10 upon *UBQLN* deficiency (Figure 1F). While gag-pol protein was more abundant in TKO cells compared to WT cells, PEG10 lacking the PPR failed to accumulate (Figure 1F-H, Figure S1C). These results identify the PEG10 PPR as a necessary region for *UBQLN2*-dependent restriction.

### Evolution of *UBQLN2*-mediated PEG10 restriction

*UBQLN2* is also unique as being the most recent gene duplication event of the *UBQLN* family. The *UBQLN2* gene is only found in eutherian mammals, commonly referred to as ‘placental’ mammals (Figure 2A) (Marín, 2014); similarly, the *PEG10* family of retrotransposon genes inserted into the mammalian genome just before the split of marsupials and eutherians (Figure 2A) (Brandt et al., 2005). While marsupial and eutherian *PEG10* share homology throughout most of the gag-pol sequence, marsupials lack the PPR at the C-terminus of pol (Figure 2A, Figure S2A). To determine the contribution of this domain to *UBQLN2*-mediated PEG10 restriction throughout mammalian evolution, cells were transfected with PEG10 fusion constructs from a variety of eutherian mammals and one marsupial, *Phascolarctos cinereus* (Koala, Figure 2B). Every PEG10 construct from eutherian mammals accumulated in *UBQLN*-deficient TKO cells (with a TKO/WT value >1), indicating reliance on UBQLNs for degradation (Figure 2C, Figure S2B). Koalas do not have a *UBQLN2* gene, and unlike the eutherian mammals tested, Koala PEG10 did not depend on UBQLN expression for its regulation. However, when the human PPR was appended to the C-terminus of Koala PEG10, its overall abundance decreased and it became more dependent on UBQLNs for restriction (Figure 2C, Figure S2B). Therefore, the PPR of PEG10, which is unique to eutherian mammals, is necessary for its relationship with UBQLN2 and confers some dependence on UBQLN2 for degradation. Together, these findings support a model whereby *UBQLN2* and *PEG10* have co-evolved a regulatory relationship involving the unique properties of UBQLN2 and the PPR of PEG10.

**Figure 2:**
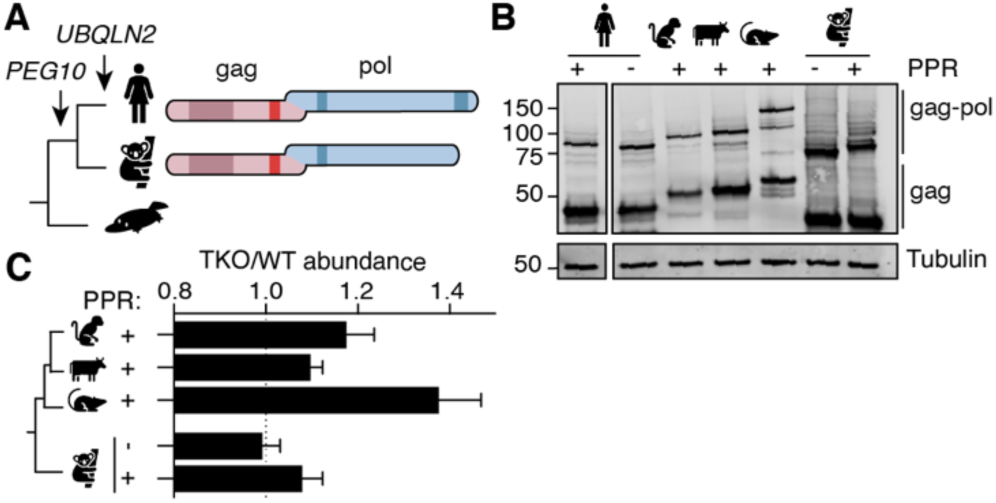
Phylogenetic investigation of the UBQLN2-PEG10 relationship. **(A)** Evolutionary schematic of PEG10 protein in eutherian and marsupial mammals. Monotremes (bottom) do not contain *PEG10* or *UBQLN2* genes. *PEG10* and *UBQLN2* appearance are highlighted with arrows. PEG10 schematic highlights the lack of C-terminal polyproline repeat region in marsupials. **(B)** Western blot demonstrating expression of mammalian PEG10 gag and gag-pol in cells. PEG10 was detected by N-terminal HA-tag. **(C)** WT or TKO cells were transfected with PEG10 reporter constructs from mammalian species and abundance of PEG10 was compared between HEK lines. Koala PEG10 lacks a PPR and was not dependent on *UBQLN* expression. Addition of the human PPR to the C-terminus of Koala PEG10 made it more dependent on *UBQLN*s for degradation. n = 4 independent experiments. For all experiments, mean ± SEM is shown.

### Human PEG10 gag-pol self-processes like a retrotransposon

The highly specific regulation of gag-pol by UBQLN2 led us to examine the unique properties of this protein in more depth. The pol region of PEG10 contains a retroviral aspartic protease domain with a classic ‘DSG’ active site motif (Figure 1D) (Clark et al., 2007), which in the ancestral Ty3 retrotransposon results in self-cleavage of capsid (CA) and nucleocapsid (NC) protein fragments with distinct functions (Clemens et al., 2011; Kirchner and Sandmeyer, 1993; Larsen et al., 2008; Sandmeyer and Clemens, 2010). Therefore, we explored the hypothesis that PEG10 gag-pol was capable of self-cleavage. Transfection with an HA-tagged form of PEG10 showed that in addition to the expected gag and gag-pol bands, there were two HA-positive lower molecular weight bands, which we hypothesized were products of self-cleavage (Figure 3A). When the active site aspartate of the PEG10 protease was mutated to alanine to disrupt proteolytic activity (gag-pol^ASG^), we observed a total disappearance of the lower molecular weight HA-tagged bands (Figure 3A), indicating that the protein products were dependent on PEG10 protease activity. Detailed biochemical analysis and bioinformatic prediction were then performed to identify the precise sites of PEG10 self-cleavage (Figure S3A-D). Together, the results suggested that PEG10 cleaves itself in two locations: AA114-115, and AA260-261. The first cleavage halves the capsid region, and the second generates a zinc-finger containing fragment reminiscent of retrotransposon and retroviral nucleocapsids (Figure 3B).

**Figure 3:**
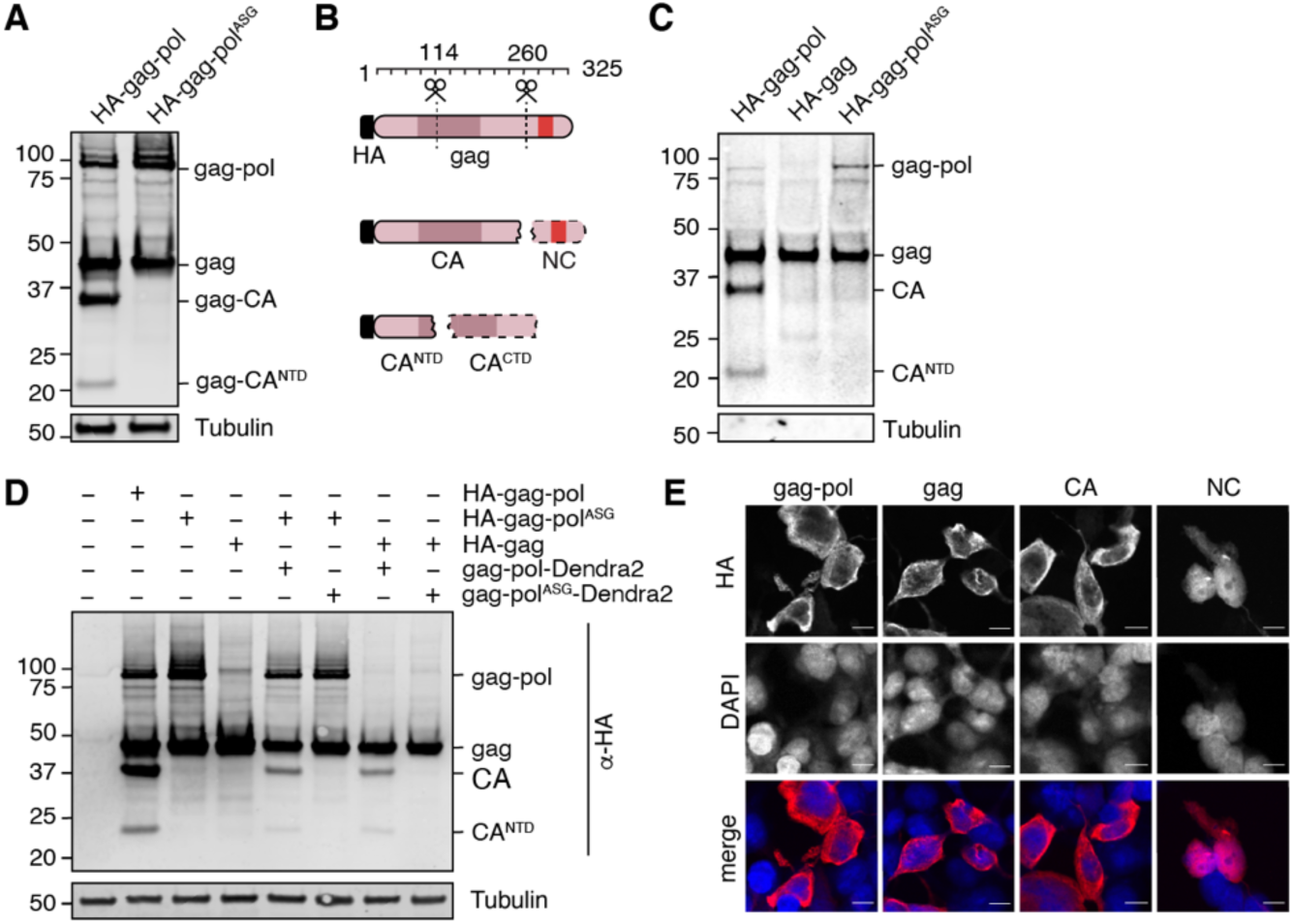
PEG10 self-cleaves to generate fragments with unique localization. **(A)** Mutation of the active site aspartic acid in the protease domain results in disappearance of cleaved PEG10 products. N-terminally HA-tagged PEG10 was expressed either as WT or ‘ASG’ protease mutant in cells and probed by western blot for HA. Based on estimated molecular weight, the fragments are estimated to encompass gag-capsid (CA) and gag- capsid(NTD) (CA^NTD^) fragments. n = 4 independent experiments. **(B)** Model of PEG10 self-cleavage. PEG10 cleaves gag to generate a liberated nucleocapsid (NC) fragment. PEG10 also cleaves the gag-CA domain into CA^NTD^ and CA^CTD^. Dotted lines indicate that the fragments are not visible by western blot due to the absence of the N-terminal HA tag. **(C)** Presence of cleaved PEG10 products in virus-like particles (VLPs). VLPs were isolated from PEG10-transfected HEK cells by ultracentrifugation and probed for cleavage products by western blot. Tubulin was used as a control for contamination of conditioned medium with cell fragments. n = 3 independent experiments. **(D)** PEG10 gag is capable of being cleaved by PEG10 gag-pol in *trans*. HA-tagged and PEG10-Dendra2 fusion constructs were co-transfected into cells and the presence of HA-tagged cleavage products was assessed by western blot. n = 2 independent experiments. **(E)** Localization of PEG10 fragments. Cells were transfected with HA-tagged PEG10 constructs and imaged by confocal microscopy after staining with an HA antibody and DAPI. Scale bar 10 μm. Shown are representative cells from 10 fields of view of each construct. n = 3 independent experiments.

Traditionally, retrotransposon and retrovirus gag-pol self-cleavage is necessary to complete the viral lifecycle. For example, proteolytic liberation of retrotransposon nucleocapsid from gag is necessary for proper capsid or virus-like particle (VLP) assembly (Larsen et al., 2008; Sandmeyer and Clemens, 2010). The PEG10 gag protein has been shown to form VLPs (Abed et al., 2019; Segel et al., 2021) that resemble those formed by retrotransposons and the gag-like gene *Arc/Arg3.1* (Ashley et al., 2018; Pastuzyn et al., 2018); therefore, we hypothesized that PEG10 self-cleavage may be necessary for proper VLP formation and release. PEG10 was overexpressed in cells and VLPs were harvested from the cultured supernatant by ultracentrifugation. Abundance of VLPs was then probed by western blot. Unlike traditional retrotransposons, self-cleavage was not a prerequisite for PEG10 VLP formation, as gag and gag-pol^ASG^ were capable of releasing VLPs with similar efficiency (Figure 3C, Figure S3E). However, like its retrotransposon ancestors, PEG10 gag-pol was capable of cleaving PEG10 gag in *trans* (Figure 3D), which suggested that a large pool of proteolytic products of gag could be generated from gag-pol activity.

Proteolytic self-processing enables novel functions for domains found in the gag and gag-pol polyproteins. For the Ty3 retrotransposon, the liberation of nucleocapsid from gag regulates the localization of capsid assembly (Larsen et al., 2008). We hypothesized that liberated PEG10 nucleocapsid may have similarly unique localization and function following self-cleavage. To test this, individual PEG10 cleavage products were expressed in cells and their localization was examined by confocal microscopy. All PEG10 proteins (gag, gag-pol, CA, and nucleocapsid) were similarly expressed and localized to the cytoplasm. Intriguingly, however, only nucleocapsid was also observed in the nucleus (Figure 3E). These data suggest that self-processing of PEG10 may reveal novel functions of its proteolytic products.

### PEG10 nucleocapsid induces changes in gene expression

The nucleocapsid fragment contains a retroviral CCHC-type zinc finger that has been reported to bind DNA (Steplewski et al., 1998) as well as RNA (Abed et al., 2019; Segel et al., 2021). This, paired with the movement of liberated nucleocapsid to the nucleus, raised the possibility that PEG10 self-cleavage may induce unique transcriptional changes. To test this hypothesis, HEK cells were transfected with either PEG10 gag-pol, gag, or nucleocapsid, and changes in gene expression were analyzed by RNA-Seq. Transfection with PEG10 gag-pol induced the most gene expression changes, followed by nucleocapsid, with gag-transfected cells showing the fewest changes compared to control (Table S1). Cluster profiling identified distinct groups of genes differentially regulated by specific PEG10 constructs. The first two groups consisted of genes that changed upon any type of PEG10 overexpression (Figure 4A), suggesting generalized responses to virus-like protein expression. One example of a gene upregulated by all forms of PEG10 expression was *TXNIP*, a regulator of oxidative stress, which is also elevated in multiple neurodegenerative conditions (Tsubaki et al., 2020) (Figure 4B). The largest cluster profile (Group 3) consisted of genes upregulated upon gag-pol and nucleocapsid transfection, but not gag transfection, highlighting the ability of the small nucleocapsid fragment to induce transcriptional changes in a manner similar to full-length gag-pol protein (Figure 4A, Figure S4A).

**Figure 4:**
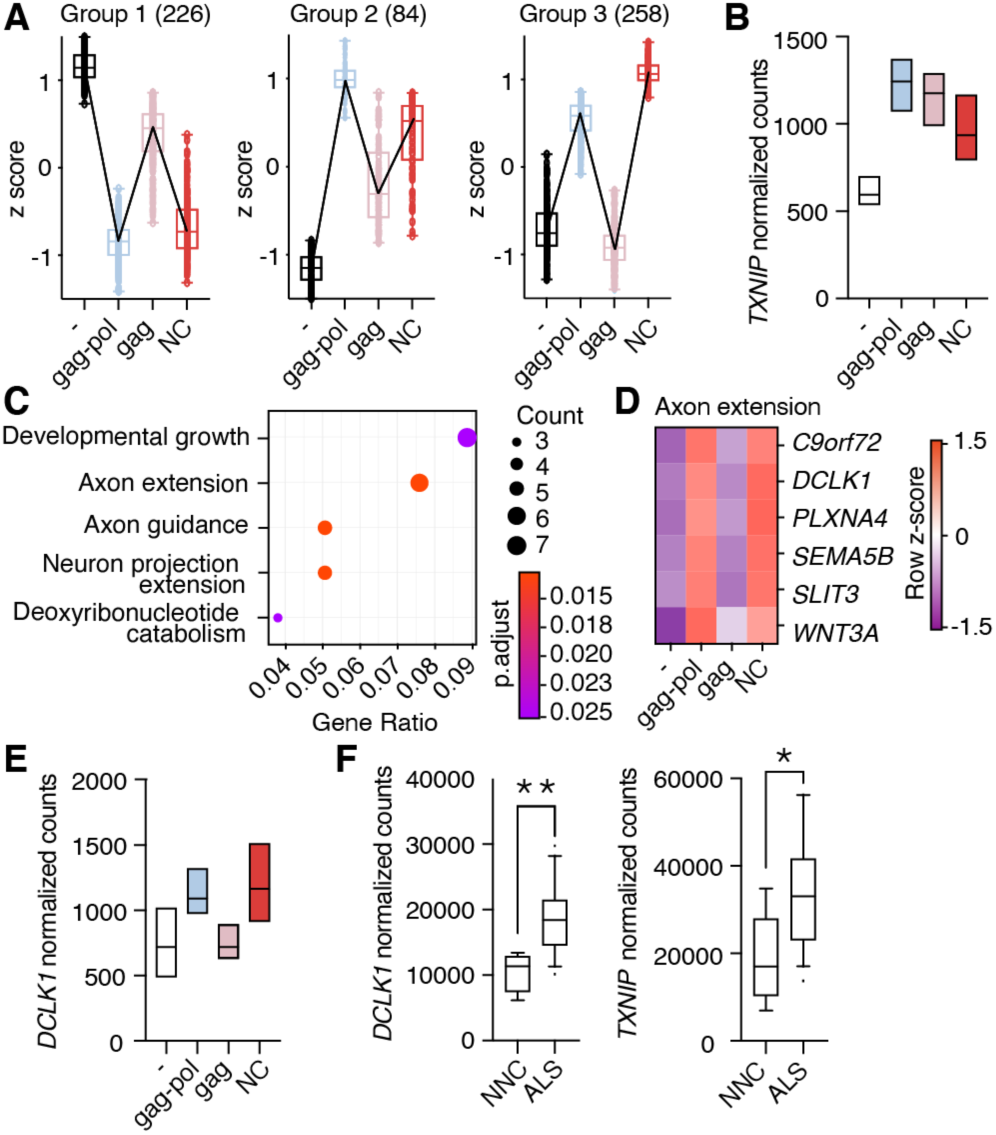
Liberated nucleocapsid alters transcription of axon extension genes. **(A)** Cluster profiling of gene expression effects as measured by RNA-Seq analysis upon PEG10 construct overexpression. The number of genes in each group is listed in parentheses. Data are shown as box and whiskers min to max with line at median. **(B)** Normalized counts of *TXNIP* transcript from RNA-Seq analysis of three biological replicates from PEG10 transfected or control transfected cells. **(C)** Top gene expression changes in NC-transfected cells by GO-term enrichment analysis. The top five GO-terms ranked by adjusted p value are shown. Adjusted p value is shown by color, and size of datapoint reflects the number of genes enriched in the pathway. **(D)** Heatmap of genes from the Axon extension GO-term showing Row z-score for each gene in the pathway. **(E)** Normalized counts of *DCLK1* from RNA-Seq analysis of PEG10-transfected cells. **(F)** RNA-Seq data from the Target ALS dataset of post-mortem lumbar spinal cords were analyzed for *DCLK1* (left) and *TXNIP* (right) counts. NNC = non-neurological control. For (B,E), data are shown as min-max floating bars with line at mean. For (F), data are shown as 5-95% box and whisker plot and significance was determined by DESeq2.

Pathway analysis of differentially expressed genes also underscored the similarities in gene regulation between gag-pol and nucleocapsid expression. Gag-pol expression resulted in an overrepresentation of pathways including female-specific sex characteristics, consistent with a role of PEG10 in placental development (Abed et al., 2019; Ono et al., 2006), as well as those involved in axon extension and remodeling (Figure S4B). Nucleocapsid expression resulted in an even stronger overrepresentation of neuronal pathways, especially pathways involved in axon guidance and extension (Figure 4C). One notable example of an axon remodeling gene was *DCLK1*, which was significantly elevated in nucleocapsid and gag-pol, but unchanged in gag-expressing cells (Figure 4D-E, Table S1). Gag expression resulted in fewer transcript changes and did not alter neuronal gene expression, highlighting the unique effects of gag-pol and nucleocapsid (Figure S4C). To better understand the transcriptional effects of PEG10 overexpression, splicing differences were examined across gag-pol, gag, and NC expression conditions. Consistent with changes to transcript abundance, both nucleocapsid and gag-pol expression resulted in splicing alteration of 150-200 genes, whereas gag had fewer effects (Table S1). There were no changes to global patterns of transcript splicing upon nucleocapsid expression (Figure S4D), nor were there global changes to mRNA trafficking (Figure S4E-F), indicating that the changes to transcriptional abundance are gene-specific.

Our data suggested a direct link between PEG10 abundance and changes to neuronal gene expression. To explore whether these transcript changes may be playing a role in human disease, we analyzed transcriptional data from post-mortem sALS patient spinal cord samples and observed similarly elevated levels of *TXNIP* and *DCLK1* transcripts (Figure 4F), suggesting that ALS involves similar pathways of transcriptional disturbance.

### PEG10 gag-pol protein is elevated in human ALS tissues

In ALS, *UBQLN2* may be dysfunctional in multiple ways. The first is through genetic mutation, which is observed in *UBQLN2*-mediated fALS (Deng et al., 2011; Williams et al., 2012) and is thought to cause both a loss of degradative function (Chang and Monteiro, 2015; Le et al., 2016) as well as a toxic gain of function by promoting misfolded UBQLN2 self-assembly (Dao et al., 2019; Sharkey et al., 2018, 2020). To test the ability of mutant *UBQLN2* to restrain PEG10 gag-pol levels, TKO cells were complemented with either WT *UBQLN2* or two known ALS-causing *UBQLN2* missense mutant alleles (Deng et al., 2011) and endogenous PEG10 was quantified by western blot (Figure 5). TKO cells expressing a WT *UBQLN2* construct were able to restrain PEG10 gag-pol to the level seen in WT cells (Figure 5), demonstrating the ability of UBQLN2 to sufficiently regulate gag-pol protein. Consistent with a loss of function phenotype, mutant *UBQLN2^P506T^*-expressing cells failed to restrict PEG10 gag-pol levels (Figure 5), whereas *UBQLN2^P497H^* had no effect on gag-pol. In all cases, gag levels were not dramatically elevated by mutant UBQLN2 expression (Figure 5C).

**Figure 5:**
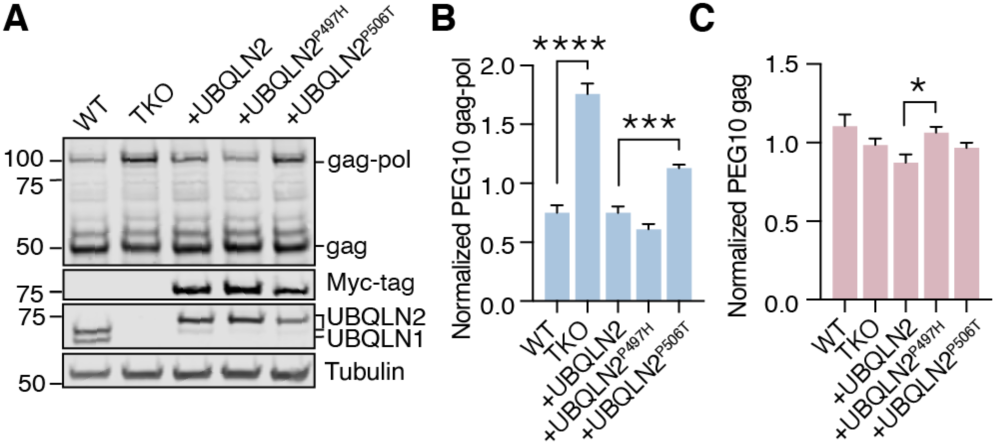
ALS-causing UBQLN2 mutation can influence PEG10 gag-pol levels. **(A)** WT or TKO cells stably transfected with doxycycline- (dox-) inducible constructs expressing *Myc-UBQLN2*, *Myc-UBQLN2^P497H^,* or *Myc-UBQLN2^P506T^* were probed for endogenous PEG10. **(B)** Quantitation of gag-pol abundance in mutant *UBQLN2*- expressing cells. Gag-pol was normalized to Tubulin and to the average intensity of each experiment. Mean ± SEM is shown for each condition. n = 5 wells per condition collected from two different passages. **(C)** Quantitation of gag abundance. Shown is mean ± SEM of n=5 wells. Multiple comparison tests were run with Bonferroni correction to compare WT and TKO cells as well as WT OE with the two mutant lines.

UBQLN2 can also be dysfunctional in the absence of overt mutation due to incorporation into ALS-associated protein aggregates (Deng et al., 2011; Williams et al., 2012), where its ability to facilitate proteasomal degradation of PEG10 may be impaired. To broadly examine the abundance of PEG10 in human patients, post-mortem lumbar spinal cord was obtained for global proteomic analysis using a tandem mass tagging (TMT) approach for liquid-chromatography mass spectrometry (LC-MS). Samples were generated from two healthy controls, six sALS cases, and one *UBQLN2^P497H^*-mediated fALS case. PEG10 is expressed at very low levels in spinal cord lysate, and in our experience is often below the technical limit of detection by LC-MS. Therefore, the samples were supplemented with a carrier channel designed to drive detection of PEG10 peptides despite the unbiased LC-MS approach (Figure 6A). Overall, 7,465 unique proteins were quantified across our samples. We observed changes in ALS tissues consistent with previously identified synaptic biomarkers of ALS and neurodegeneration, such as a significant reduction of neurogranin (Kvartsberg et al., 2019; Vijayakumar et al., 2019) (Figure S5A-C, Table S2). Seven peptides were identified from the gag region of PEG10, and three peptides were identified from the pol region (Figure S6A-B). While gag was not changed in *UBQLN2*-mediated or sporadic ALS samples (Figure 6B), peptides originating from PEG10 pol were significantly enriched in ALS compared to healthy controls (Figure 6C). This was specific to the protein level, as *PEG10* transcript counts were not elevated in ALS (Figure S6C). When PEG10 gag and pol were considered as unique proteins (2 peptides per protein minimum, Table S2), PEG10 pol was among the most upregulated proteins in all ALS cases compared to healthy controls (Figure 6D). Its abundance was also associated with a decrease in neurogranin and neurofilament medium (Figure S6D-E), highlighting the potential of PEG10 pol as a novel biomarker for ALS. Taken together, the accumulation of PEG10 gag-pol in ALS tissue, paired with our findings of PEG10 self-cleavage and effects on gene expression, suggest that this pathway may represent a novel pathological contribution to the development of ALS.

**Figure 6:**
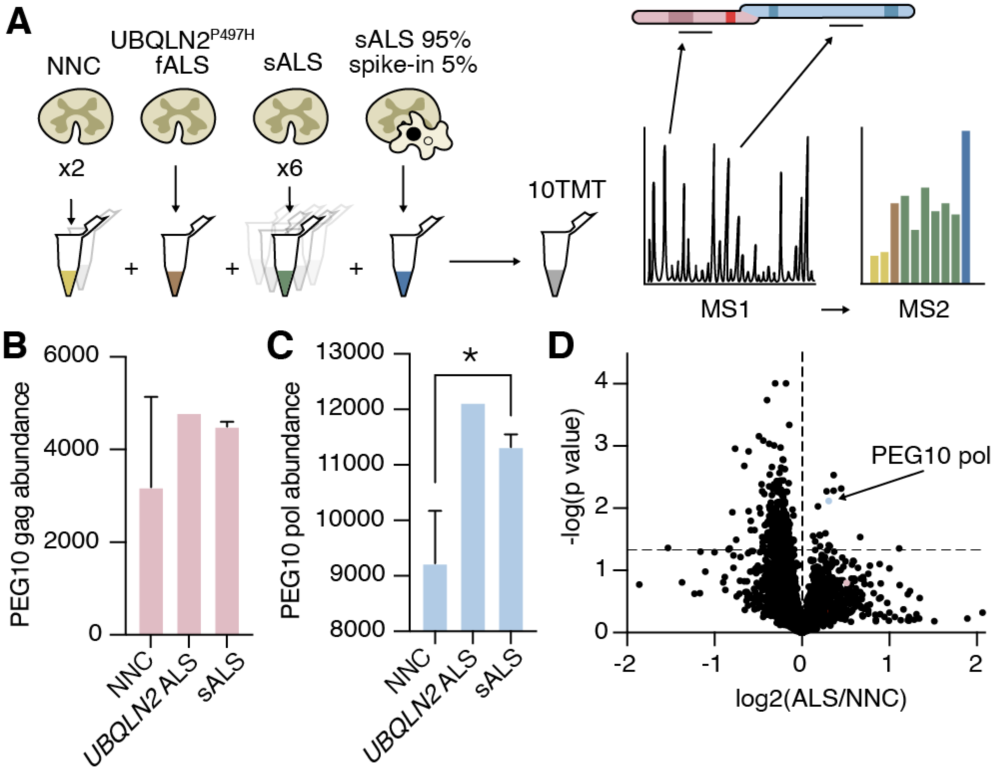
PEG10 gag-pol protein accumulates in human ALS. **(A)** Schematic illustrating multiplexed global proteomic strategy to quantify PEG10 protein from human lumbar spinal cord. Two non-neurological controls (NNC), one fALS case with a *UBQLN2* mutation, and six sporadic ALS cases were combined with a ‘spike-in’ PEG10 channel containing 5% lysate from cells transfected with HA-PEG10 gag-pol and 95% spinal cord lysate to normalize proteomic background complexity. All ten samples were labeled with tandem mass tags (TMT) and run as a 10-plex on LC-MS2. **(B-C)** Abundance of PEG10 gag (B), and pol (C), in human spinal cord. Significance was determined by Student’s t-test. **(D)** Global proteomic analysis with 7,465 individual proteins quantified. All ALS samples were grouped together, and two non-neurological controls (NNC) were grouped to generated log_2_ ratio of protein abundance and significance calculation by homoscedastic unpaired t-test. PEG10 pol is highlighted in blue, and PEG10 gag (not significant) is highlighted in pink.

## Discussion

Here, we have explored in detail the regulatory mechanisms and cellular consequences of PEG10 gag-pol self-processing and observed the accumulation of PEG10 gag-pol in both sporadic and *UBQLN2*-mediated ALS. PEG10 gag-pol exhibited protease-dependent self-cleavage and generated a nuclear-bound fragment that is reported to bind nucleic acids (Abed et al., 2019; Segel et al., 2021; Steplewski et al., 1998). This fragment was sufficient to change transcript abundance in the cell, including the expression of genes involved in axon remodeling, linking PEG10 dysregulation to neuronal dysfunction. Finally, we observed a specific accumulation of PEG10 gag-pol in spinal cord tissue of ALS patients, suggesting that PEG10 activity may contribute to disease in humans.

Data outlined here suggest a specific regulatory and evolutionary relationship between UBQLN2 and the domesticated retrotransposon, PEG10. The regulatory relationship between UBQLN2 and PEG10 is specific in two ways: first, UBQLN2 is the only UBQLN capable of restraining PEG10 abundance. The unique ability of UBQLN2 to mediate PEG10 degradation is likely dependent on the PXX domain of UBQLN2, which is not found in other *UBQLN* genes, and is a mutational hotspot in *UBQLN2*-mediated fALS (Deng et al., 2011). Rescue of UBQLN TKO cells with *UBQLN2^P506T^* failed to reduce PEG10 gag-pol levels to those of WT cells, which is consistent with our hypothesis. Second, only the gag-pol form of PEG10 is a client of UBQLN2, due to the presence of the pol domain’s unique PPR. It is notable that both protein domains that are necessary for the UBQLN2-PEG10 relationship exist only in eutherian mammals, suggesting an evolutionary relationship. In comparison, the rest of the *UBQLN* gene family is more conserved among eukaryotes: *UBQLN3*, *5*, and *L* are shared among earlier mammalian ancestors, *UBQLN1* first appears in fish species, and *UBQLN4* is thought to reflect adaptation of the ancestral *Dsk2* gene in single-celled eukaryotes (Marín, 2014). These points, paired with the finding that PEG10 was by far the most dysregulated protein in global proteomic studies of *UBQLN2* loss (Whiteley et al., 2021), raise the intriguing possibility that contrary to its presumed role as a regulator of generalized protein degradation, *UBQLN2* may have evolutionarily arisen specifically to restrain PEG10 abundance. Further, the unique relationship between *PEG10* and *UBQLN2* highlights a potential genetic conflict between PEG10 expression requirements in placental development (Ono et al., 2006) with pathological roles in neural tissue which would have necessitated the evolutionary development of a tissue-specific inhibitor of gag-pol activities. PEG10 has recently emerged as a potential player in the neurodevelopmental disorder Angelman’s syndrome, where its abundance was elevated in disease (Pandya et al., 2021). This observed dysregulation of PEG10, as well as the unique enrichment of UBQLN2 in neuromuscular tissues, further supports our hypothesis.

Our work also adds insight into the biology of PEG10. PEG10 self-cleavage is reminiscent of the self-processing of retrotransposons and resulted in unique transcriptional changes to the cell. Neuronal genes were some of the most altered transcripts upon gag-pol and nucleocapsid transfection, implicating a link between protease-dependent PEG10 self-cleavage, nucleocapsid liberation, and neuronal dysfunction. However, further work is needed to identify the molecular mechanism by which nucleocapsid mediates transcriptional changes. The nucleocapsid fragment contains a classic retroviral CCHC-type ‘zinc finger’, and PEG10 has been reported to bind cellular mRNAs (Abed et al., 2019; Segel et al., 2021), as well as DNA (Steplewski et al., 1998). Therefore, nucleocapsid may behave as a classical transcription factor, or as a regulator of transcript abundance through other means such as regulation of gene-specific mRNA stability, trafficking, or splicing. These possibilities remain to be elucidated. Our work supports an important role for PEG10 gag-pol accumulation as a mechanism of ALS disease progression. Even in the absence of overt *UBQLN2* mutation, PEG10 gag-pol protein is accumulated in post-mortem tissue of sALS patients compared to non-neurological controls. We hypothesize that this disease-wide accumulation is related to the inclusion of UBQLN2 in sALS-associated protein aggregates (Deng et al., 2011; Williams et al., 2012), which would lead to a similar loss of function phenotype. Only PEG10 pol peptides were upregulated in spinal cord samples of both fALS and sALS, indicating a specific upregulation of protease-containing gag-pol PEG10 in disease. Strikingly, among the more than 7,000 proteins quantified by our analysis, PEG10 pol was the fifth most significantly upregulated protein in spinal cord tissue of ALS patients, suggesting a strong connection to disease and the potential utility of PEG10 as a novel biomarker for ALS progression. This further suggests that PEG10 gag-pol accumulation may be considered alongside TDP43, FUS, and UBQLN2 mislocalization and aggregation as shared hallmarks of familial and sporadic ALS (Blokhuis et al., 2013). Ultimately, further understanding of PEG10 biology in the context of ALS may provide novel avenues for therapeutic development.

## Acknowledgements

We would like to acknowledge the Target ALS Human Postmortem Tissue Core, New York Genome Center for Genomics of Neurodegenerative Disease, Amyotrophic Lateral Sclerosis Association and TOW Foundation for post-mortem tissue samples and RNA-Seq reads. We would also like to acknowledge The Shared Instruments Pool of the Department of Biochemistry at CU Boulder, the Biochemistry Cell Culture Core Facility, the Biochemistry Flow Cytometry Core Facility, and the BioFrontiers Advanced Light Microscopy Core for their assistance with shared equipment. Drs. Miguel Prado and Ramanujan Hegde provided WT, TKO, and *UBQLN2*-rescue HEK293 cells for experiments. We thank Dr. Yoseph Barash and members of his lab for providing us early access to the MAJIQ classifier and for assistance in classifier setup. We would also like to thank Drs. Edward Chuong, Roy Parker, Aaron Holling, and Aaron Whiteley for critical reading of our manuscript.

## Author Contributions

H.H.B performed and analyzed RNA-Seq, western blotting, and microscopy. J.E.R. performed and analyzed ES cell blotting and phylogenetic flow cytometry data. S.N.L. performed and analyzed mutant UBQLN2 HEK293 cell data and global proteomic data, and performed statistical analysis. W.C. performed and analyzed VLP preparations and blotting. C.C.E. advised on proteomic study design and performed global proteomics. C.I.L. performed ES cell blotting. A.M.W. conceived and designed the study, analyzed and interpreted data, and wrote the manuscript. All authors edited the manuscript and support the conclusions.

## STAR Methods

### Antibodies Used

| Immunogen | Clonality | Species | Vendor | Catalog # | Usage | Purpose |
| --- | --- | --- | --- | --- | --- | --- |
| UBQLN1/2 | Monoclonal M03 | Mouse | Abnova | H00029978 | 1:1000 | WB |
| UBQLN4 | Polyclonal | Rabbit | GeneTex | GTX85267 | 1:1000 | WB |
| Myc tag | Monoclonal GT002 | Mouse | Sigma | SAB270219<br>2 | 1:2000 | WB |
| PEG10 | Polyclonal | Rabbit | Proteintech | 14412-1-AP | 1:1000 | WB |
| HA | Monoclonal HA-7 | Mouse | Sigma | H3663 | 1:1000 | WB, IF |
| Tubulin | Monoclonal DM1A | Mouse | Novus | NB100-690 | 1:20000 | WB |
| α-mouse IgG 680 | Polyclonal | Goat | Licor | 926-68070 | 1:20000 | WB |
| α-mouse IgG 800 | Polyclonal | Goat | Licor | 926-32210 | 1:20000 | WB |
| α-rabbit IgG 680 | Polyclonal | Goat | Licor | 926-68071 | 1:20000 | WB |
| α-rabbit IgG 800 | Polyclonal | Goat | Licor | 926-32211 | 1:20000 | WB |
| Alexa568 α-mouse | Polyclonal | Goat | Invitrogen | A11004 | 1:300 | IF |

### Cell lines

WT HEK293 cells and HEK cells lacking *UBQLNs* 1,2, and 4 (‘TKO’) were a gift from Dr. Ramanujan Hegde of the Medical Research Council Laboratory of Molecular Biology. WT and TKO HEK293 cells were cultured in Dulbecco’s modified Eagle’s medium (Invitrogen) supplemented with 1% penicillin/streptomycin (Invitrogen), 1% L-glutamine (R&D Systems, Inc.), and 10% FBS (Millipore Sigma). HEK293 TKO cells stably transfected with doxycycline- (dox-) inducible constructs expressing *Myc-UBQLN2*, or *Myc- UBQLN2^P497H^* or *Myc-UBQLN2^P506T^* were provided by Dr. Hegde via Dr. Miguel Prado and are described in Itakura et al. Dox-inducible cells were cultured in DMEM supplemented with 1% penicillin/streptomycin (Invitrogen), 1% L-glutamine (R&D Systems, Inc.), and 10% tet-system approved FBS (Gibco), as contaminating doxycycline in standard FBS was sufficient to stimulate high levels of *UBQLN2* expression. Leaky expression of *Myc- UBQLN2* constructs in dox-free medium resulted in endogenous levels of tagged UBQLN2 expression.

Human H9 hESCs lacking either *UBQLN1*, *2*, or *4*, were generated at the Harvard Medical School Cell Biology Initiative for Genome Editing and Neurodegeneration according to(Whiteley et al., 2021). hESCs were cultured in either E8 (StemCell Technologies) or TeSR-E8 medium (StemCell Technologies) on 6-well tissue culture plates with Matrigel (Corning, lot #0048006). Medium was changed daily. Cells were passaged by treatment in 0.5 mM EDTA (Sigma Aldrich) in sterile PBS (Invitrogen) and replated in media at an approximate 1:6 dilution. The remaining non-passaged cells were washed in D-PBS three times and pelleted at 300 x g for analysis.

### Cell Transfection

WT or TKO HEK293 cells were grown to 70% confluency in 12-well plates and transfected with 1 μg plasmid DNA in Lipofectamine 2000 (Invitrogen) and Opti-Mem medium (Invitrogen), according to manufacturer’s instructions. After 48 hours, cells were harvested for western blot, qPCR, RNA-Seq, or immunofluorescence.

### Human Tissue Samples

Human tissue samples were acquired from the Target ALS Multicenter Human Postmortem Tissue Core. Unfixed, full-thickness sections of lumbar spinal cord were obtained from two non-neurological controls, one ALS patient with a pathogenic *UBQLN2* mutation, and seven sporadic ALS cases. All cases were from females. More detail can be found in Supplementary Information Table 2.

### Cloning

All constructs were designed using Gibson or restriction cloning (see list of constructs) and transformed into chemically competent DH5ɑ *E. coli* cells (Invitrogen). Transformed *E. coli* were plated on either 50 μg/mL kanamycin (Teknova) or 100 μg/mL carbenicillin (Gold Biotechnology) LB agar (Teknova) plates overnight at 37° C. Single colonies were picked and grown overnight in 5 mL LB Broth (Alfa Aesar) with kanamycin or carbenicillin at 37°C with shaking at 220 rpm. The following day, shaking cultures were mini-prepped (Zymo) and sent for Sanger Sequencing. Sequence verified samples were then grown in 50 mL LB Broth overnight with appropriate antibiotic at 37° C with shaking at 220 rpm. 50 mL cultures were midi-prepped (Zymo) for transfection.

### Constructs

| Construct Name | Insert | Species | NCBI protein or transcript reference | Vector |
| --- | --- | --- | --- | --- |
| - | LIFEACT (Riedl et al., 2008) |  | n/a | pCDNA3.1 |
| gag-pol | PEG10 (AA1-708) | *Homo sapiens* | NP_055883.2 | pCDNA3.1 |
| gag-pol<sup>ASG</sup> | PEG10 (AA1-708)* D370A | *Homo sapiens* | NP_055883.2 | pCDNA3.1 |
| gag | PEG10 (AA1-325) | *Homo sapiens* | NP_001035242.1 | pCDNA3.1 |
| CA | PEG10 (AA1-259) | *Homo sapiens* | NP_001035242.1 | pCDNA3.1 |
| NC | PEG10 (AA260-325) | *Homo sapiens* | NP_001035242.1 | pCDNA3.1 |
| gag111 | PEG10(AA1-111) | *Homo sapiens* | NP_001035242.1 | pCDNA3.1 |
| gag-pol Dendra2 | PEG10 (AA1-708)-Dendra2 | *Homo sapiens* | NP_055883.2 | pDendra2 |
| gag-pol<sup>ASG</sup> Dendra2 | PEG10 (AA1-708)*<br>D370A-Dendra2 | *Homo sapiens* | NP_055883.2 | pDendra2 |
| ΔPPR Dendra2 | PEG10 (AA1-681)-<br>Dendra2 | *Homo sapiens* | NP_055883.2 | pDendra2 |
| *Macaca Mulatta*<br>gag-pol Dendra2 | PEG10<br>(AA1-742)-Dendra2 | *Macaca mulatta* | NP_001165893.2 | pDendra2 |
| *Bos taurus* gag-pol<br>Dendra2 | PEG10<br>(AA1-788)-Dendra2 | *Bos taurus* | NP_001120682.1 | pDendra2 |
| *Mus musculus* gag-pol Dendra2 | PEG10<br>(AA1-1006)-Dendra2 | *Mus musculus* | NP_570947.2 | pDendra2 |
| *Phascolarctos cinereus* gag-pol<br>Dendra2 | PEG10<br>(AA1-624)-Dendra2 | *Phascolarctos cinereus* | XM_021000084.1<br>(+ downstream CDS<br>continuation to *pol*<br>stop codon) | pDendra2 |
| *Phascolarctos cinereus* gag-pol<br>Dendra2 + HsPPR | Pc PEG10 (AA1-624)<br>+<br>Hs PEG10 (AA682-708)<br>Dendra2 | *Phascolarctos cinereus* and<br>*Homo sapiens* | XM_021000084.1<br>(+ downstream CDS<br>continuation to *pol*<br>stop codon)<br>And<br>NP_055883.2 | pDendra2 |

### Flow Cytometry

WT and TKO HEK293 cells were transfected in 96-well plates. Cells were harvested 48 hours after transfection in FACS Buffer (D-PBS, 2% FBS, 0.1% Sodium Azide) and analyzed on a BD FACSCelesta. Triplicate wells were transfected within a plate to serve as technical replicates, and experiments were performed four independent times. FlowJo software was used for data analysis.

Cells were first gated in the FSC-A vs. SSC-A using the polygon gating tool. Within the ‘cells’ population, CFP positive cells were gated on 405 nm vs. SSC-A. The Dendra Green/CFP parameter was created by deriving a novel parameter of the 488 reference by the 405 reference, and making a logarithmic scale with a minimum of 0.0001 and a maximum of 10. The geometric mean of the custom Dendra Green/CFP parameter from the CFP positive population was exported and used to generate graphs.

### Western Blotting

Cell pellets were centrifuged, washed in PBS and lysed in urea buffer (8 M urea, 75 mM NaCl, 50 mM HEPES pH 8.5, 1x tab cOmplete Mini EDTA-free protease inhibitor cocktail tablet (Sigma Aldrich)). Lysis was performed by vortex and incubation for 15 minutes at room temperature. Lysate was centrifuged for 10 minutes at 21,300 x g and the supernatant was collected as sample.

Protein was quantified by BCA (Pierce) and 1x Laemmli sample buffer supplemented with βME (Sigma Aldrich) was added to samples before SDS-PAGE. Samples were run in NuPage MES Running Buffer (Invitrogen) on a 4 to 12% NuPage Bis-Tris gel (Invitrogen) and wet transferred on nitrocellulose membrane (Amersham Protran) for either 90 minutes at 100 V on ice (BioRad Mini-Blot Transfer) or 60 minutes at 10 V (Invitrogen Mini Tank Blot Module).

Membranes were blocked using 1:1 LICOR blocking buffer and 1x TBS (50 mM Tris-Cl pH 7.4, 150 mM NaCl), for 30 minutes at room temperature. Membranes were incubated in primary antibody overnight at 4° C and washed in 1x TBST (1x TBS, 0.1% Tween, VWR) in three five-minute intervals. Membranes were then incubated in LICOR secondary antibody for 30 minutes in the dark. After 3x more washes, banding patterns were visualized using LICOR Odyssey CLx and data analysis was performed using LICOR ImageStudio Software. Each protein quantification was normalized to the average intensity across all samples in each replicate western blot to correct for technical variation across experiments.

### Virus-Like Particle Isolation

HEK293 cells were plated in a 6-well plate at a density of 4×10^5^ cells per well. 24 hours after plating, cells were transfected and media was replaced 6 hours later. Cultured media was harvested 48 hours after transfection and pre-cleared by centrifugation at 2000 x g for 15 minutes at 4° C. In parallel, cell lysate was collected for western blot as previously described. The VLP fraction was isolated by ultracentrifugation (Beckman Coulter) at 134,000 x g for 4 hours with a 30% sucrose cushion. After ultracentrifugation, media and sucrose were aspirated, and the VLP-containing pellet was resuspended in 8M urea lysis buffer. VLP production was analyzed by western blot.

### Phylogenetic alignment of PEG10

PEG10 protein sequences were curated from NCBI for selected eutherian mammals, and manually curated from marsupial mRNA sequences from NCBI due to a lack of automatically annotated frameshifting sites. Sequences were then aligned with Geneious 3.0 using a MUSCLE-based algorithm with 8 iterations. Alignment was visualized with unique colors for each amino acid.

### Structure prediction

Structure prediction for the PEG10 gag protein (AA1-325) was performed using the Phyre 2.0 webserver(Kelley et al., 2015) using the intensive modeling mode. 243 of 325 amino acids were modeled with >90% confidence, with amino acids 89-314 predicted with confidence >99% against reference structures including *Saccharomyces* Ty3, *Drosophila* and *Rattus* Arc, HIV, and a partial structure of *Homo sapiens* PEG10 gag. The predicted PEG10 structure was visualized using UCSF Chimera.

### Immunofluorescence

24 hours after transfection, HEK293 cells were re-plated onto Alcian Blue- (Newcomer Supply) treated round coverslips (Electron Microscopy Sciences) in 24-well plates and cultured overnight at 37° C. 24 hours later, coverslips were harvested and fixed in 4% PFA (Thermo Scientific Pierce). Cells were then either submerged in 1% PFA for overnight storage or washed three times in 1x PBS. Cells were permeabilized in 0.25% Triton-X (Sigma Aldrich) in PBS and incubated in blocking buffer (7.5 % BSA (Gibco) diluted to 5% in PBS, 0.1% tween) for 30 minutes. Cells were incubated for 1 hour in primary antibody before three 5-minute washes in 1x PBS-T (0.1% Tween in PBS). Cells were then incubated in secondary antibody for one hour in the dark. Cells underwent three more 5-minute 1x PBS-T washes and were then rinsed three times in DEPC water. After sufficient drying, 5-10 mL of Prolong Gold DAPI anti-fade mounting media (Invitrogen) was added to coverslips. Coverslips were then mounted on clear microscope slides and cured overnight in the dark at room temperature before imaging.

### Oligo dT Fluorescence *in situ* Hybridization

Transfected WT HEK293 cells were cultured on round coverslips (Electron Microscopy Sciences) and hybridized according to Stellaris protocol for hybridization of adherent cells (Biosearch Technologies). A T30 Poly A probe (Stellaris, Biosearch Technologies) was used to detect polyA mRNA tails as a measure of total mRNA by cellular compartment. 10 images were obtained for each transfection condition and randomly assigned image names for blind quantification. Distinct single cells were quantified using FIJI software XOR function to quantify mean signal intensity of nucleus and cytoplasm. Nucleus to cytoplasmic ratio was calculated for a minimum of 60 cells for each condition.

### Confocal Microscopy

Microscopy was performed on a Nikon AR1 LSM confocal microscope maintained by the BioFrontiers Advanced Light Microscopy Core using a 20x Air objective and NIS Elements Nikon software. Acquisition intensity and pinhole size were fixed across samples to control for signal intensity and variability. For visualization purposes only, image intensity of visualized channels was increased from acquisition parameters according to FIJI software parameters.

### Sample preparation for RNA Sequencing

HEK293 cells were grown in 12-well plates, transfected for overexpression of genes of interest, and collected for RNA isolation 48 hours later. Cells were pelleted and RNA was extracted using the RNEasy Mini Kit (Qiagen) with on-column DNAse digestion (Qiagen). Isolated RNA was quantified and quality controlled by nanodrop, concentration was normalized, and samples were stored at -80° C.

### RNA-Sequencing Analysis

Poly A Selected Total RNA Library paired-end sequencing was performed at Anschutz Medical Campus on an Illumina NovaSEQ 6000. Sequencing produced between 24-104 million filtered paired-end reads across all samples. Quality of reads was determined using FastQC (the average reads/base quality for all samples in the lane was at least 88%>=Q30) and reads were mapped to GRCh38.p13(Frankish et al., 2019) using STAR version 2.7.3 (Dobin et al., 2013). STAR alignment .bam files were indexed and sorted before count matrix generation using Samtools 1.8 and featureCounts software package (Li et al., 2009).

Count files were converted to readable format in unix and imported into RStudio for DESeq2 analysis using R. Data were quality controlled by estimating size factors and genewise dispersion estimates for variance in gene expression. Shrinking was used to fit dispersion curves and principal component analyses dictated design parameters for differential gene expression analysis. Gene expression patterns were tracked using DESeq2 (Love et al., 2014) using harvest date and transfection construct as major variables, as well as Cluster Profiling(Yu et al., 2012), and GO Term expression (Luo and Brouwer, 2013; Yu et al., 2015) analyses. Significance of gene expression changes was determined with a p-adjusted cutoff of .05. Gene groups were determined with DEGReport (Pantano, 2021) using a reduced cluster model in which outliers of cluster distribution were removed.

Pathway analysis was performed using the enrichGO program (Yu et al., 2015) on all GO-term pathways with a log_2_foldchange cutoff of 0.5 and a p value of 0.05 of significantly changed genes for each pairwise analysis (pCDNA negative control vs. gag-pol, vs. gag, and vs. NC). The top 5 pathways by p value were visualized.

Splicing analysis was performed using the MAJIQ Quantifier followed by the MAJIQ Builder to determine differentially spliced genes (Vaquero-Garcia et al., 2016), and visualized using the MAJIQ Voila Viewer with a Δψ threshold of 0.1 and significance of 0.05. Splice variant classification analysis was performed using the MAJIQ classifier (Vaquero-Garcia et al., 2021) with permission and assistance from Dr. Yoseph Barash.

### Target ALS dataset RNA-Seq analysis

Raw RNA-Seq reads from lumbar spinal cord of the Target ALS: New York Genome Center dataset were obtained; at the time of analysis, this dataset included 40 Classical/Typical ALS cases, 5 non-neurological controls, and multiple samples from other neurodegenerative diseases for a total of 51 samples. Approximately half of the samples were male/female. Reads were aligned to the human genome (hg38) using STAR version 2.5.2b as above (Dobin et al., 2013), and analyzed in RStudio with DESeq2 (Love et al., 2014) including sex and ‘Subject.Group.Subcategory’ (disease type) as major variables. Significance of gene expression changes was determined with a p-adjusted cutoff of .05, and normalized counts were used for visualization of target genes.

### Sample preparation for mass spectrometry analysis

Human spinal cord samples were first sectioned on a cryostat (Leica) to ensure even tissue representation of protein sample. Ten to twenty 15 µm-thickness sections from each patient were homogenized in 8M urea lysis buffer, lysate was spun at 15,000 rpm for 15 minutes at 4° C to remove insoluble material, and supernatant protein content was quantified by BCA analysis (Pierce). Separately, HEK cells were transfected with *Homo sapiens* HA-PEG10 gag-pol, lysed 48 hours later, and mixed in a 95:5 ratio of sALS spinal cord lysate to HEK cell lysate. Approximately 100-200 μg of each sample was aliquoted and delivered to the Proteomics and Mass Spectrometry Core Facility in the Department of Biochemistry at the University of Colorado, Boulder, for TMT labeling.

Human lumbar spinal cord tissue samples in 8M urea were reduced and alkylated with the addition of 5% (w/v) sodium dodecyl sulfate (SDS), 10 mM tris(2-carboxyethylphosphine) (TCEP), 40 mM 2-chloroacetamide, 50 mM Tris-HCl, pH 8.5 and incubated shaking at 1000 rpm at room temperature for 60 minutes then cleared via centrifugation at 17,000 x g for 10 minutes at 25°C. Lysates were digested using the SP3 method(Hughes et al., 2014). Briefly, 200 µg carboxylate-functionalized speedbeads (Cytiva Life Sciences) were added to approximately 100 µg protein lysate. Addition of acetonitrile to 80% (v/v) induced binding to the beads, then the beads were washed twice with 80% (v/v) ethanol and twice with 100% acetonitrile. Proteins were digested in 50 mM Tris-HCl buffer, pH 8.5, with 1 µg Lys-C/Trypsin (Promega) and incubated at 37° C overnight. Tryptic peptides were desalted using HLB Oasis 1cc (10mg) cartridges (Waters) according to the manufactures instructions and dried in a speedvac vacuum centrifuge. Approximately 30 µg of tryptic peptide from each human tissue sample was labeled with TMT 10 plex (Thermo Scientific) reagents according to the manufacturer’s instructions. The multiplexed sample was cleaned up with a HLB Oasis 1cc (10mg) cartridge. Approximately 50 µg multiplexed peptides were fractionated with high pH reversed-phase C18 UPLC using a 0.5 mm X 200 mm custom packed UChrom C18 1.8 µm 120Å (nanolcms) column with mobile phases 10mM aqueous ammonia, pH10 in water and acetonitrile (ACN). Peptides were gradient eluted at 20 µL/minute from 2 to 40% ACN in 40 minutes concatenating for 12 fractions using a Waters M-class UPLC (Waters). Peptide fractions were then dried in a speedvac vacuum centrifuge and stored at -20°C until analysis.

### Mass spectrometry analysis

High pH peptide fractions were suspended in 3% (v/v) ACN, 0.1% (v/v) trifluoroacetic acid (TFA) and approximately 1 µg tryptic peptides were directly injected onto a reversed-phase C18 1.7 µm, 130 Å, 75 mm X 250 mm M-class column (Waters), using an Ultimate 3000 nanoUPLC (Thermos Scientific). Peptides were eluted at 300 nL/minute with a gradient from 4% to 16% ACN over 120 minutes then to 25% ACN in 5 minutes and detected using a Q-Exactive HF-X mass spectrometer (Thermo Scientific). Precursor mass spectra (MS1) were acquired at a resolution of 120,000 from 350 to 1500 m/z with an automatic gain control (AGC) target of 3E6 and a maximum injection time of 50 milliseconds. Precursor peptide ion isolation width for MS2 fragment scans was 0.7 m/z with a 0.2 m/z offset, and the top 15 most intense ions were sequenced. All MS2 spectra were acquired at a resolution of 45,000 with higher energy collision dissociation (HCD) at 32% normalized collision energy. An AGC target of 1E5 and 100 milliseconds maximum injection time was used. Dynamic exclusion was set for 20 seconds with a mass tolerance of ±10 ppm. Raw files were searched against the Uniprot Human database UP000005640 downloaded November 2, 2020 using MaxQuant v.1.6.14.0. Cysteine carbamidomethylation was considered a fixed modification, while methionine oxidation and protein N-terminal acetylation were searched as variable modifications. All peptide and protein identifications were thresholded at a 1% false discovery rate (FDR).

For visualization of data, likely contaminants, reverse peptides, and proteins quantified by only one peptide were removed. P values were calculated by Student’s t-test (unpaired, homoscedastic variance) combining both non-neurological control samples and combining all ALS cases (including sporadic and *UBQLN2*-mediated).

### Statistical Analysis

For western blots, values (normalized to Tubulin and batch-corrected) were compared using the appropriate statistical test by determining normality using a Shapiro-Wilk test and variance using Bartlett’s test. For values that were normally distributed and had equal variance, a standard one-way ANOVA was first used to compare differences between the means across all groups. If all groups were not normally distributed a Kruskal-Wallis test was used. Appropriate multiple comparisons tests were utilized to determine which groups significantly varied. For normal distributions, a Bonferroni’s multiple comparisons test was used to compare means directly. For non-normally distributed results, a Dunn’s multiple comparisons test was used.

For proteomic analysis, values were compared with an unpaired t-test and a threshold of p<0.05 was used for significance. For RNA-Seq data, statistical analysis is described above using DESeq2 and adjusted p values.

For all figures, statistical tests are listed in the figure legend and *p<0.05, **p<0.01, ***p<0.001, and ****p<0.0001.

### Data Availability

RNA-Seq data is available on the Gene Expression Omnibus (GEO) at XXX. Proteomics data is available on PRIDE at XXXX.

## Data and Materials Availability

All data is available in the manuscript or the supplementary materials. Correspondence and material requests should be directed to A. M. Whiteley at.

## References

Abed, M., Verschueren, E., Budayeva, H., Liu, P., Kirkpatrick, D.S., Reja, R., Kummerfeld, S.K., Webster, J.D., Gierke, S., Reichelt, M., et al. (2019). The Gag protein PEG10 binds to RNA and regulates trophoblast stem cell lineage specification. PLoS ONE 14, e0214110.

Akamatsu, S., Wyatt, A.W., Lin, D., Lysakowski, S., Zhang, F., Kim, S., Tse, C., Wang, K., Mo, F., Haegert, A., et al. (2015). The Placental Gene PEG10 Promotes Progression of Neuroendocrine Prostate Cancer. Cell Reports 12, 922–936.

Ashley, J., Cordy, B., Lucia, D., Fradkin, L.G., Budnik, V., and Thomson, T. (2018). Retrovirus-like Gag Protein Arc1 Binds RNA and Traffics across Synaptic Boutons. Cell 172, 262–274.e11.

Blokhuis, A.M., Groen, E.J.N., Koppers, M., van den Berg, L.H., and Pasterkamp, R.J. (2013). Protein aggregation in amyotrophic lateral sclerosis. Acta Neuropathol 125, 777–794.

Brandt, J., Veith, A.M., and Volff, J.-N. (2005). A family of neofunctionalized Ty3/gypsy retrotransposon genes in mammalian genomes. Cytogenet Genome Res 110, 307–317.

Brown, R.H., and Al-Chalabi, A. (2017). Amyotrophic Lateral Sclerosis. N Engl J Med 377, 162–172.

Chang, L., and Monteiro, M.J. (2015). Defective Proteasome Delivery of Polyubiquitinated Proteins by Ubiquilin-2 Proteins Containing ALS Mutations. PLoS ONE 10, e0130162.

Clark, M.B., Jänicke, M., Gottesbühren, U., Kleffmann, T., Legge, M., Poole, E.S., and Tate, W.P. (2007). Mammalian Gene PEG10 Expresses Two Reading Frames by High Efficiency –1 Frameshifting in Embryonic-associated Tissues. J. Biol. Chem. 282, 37359–37369.

Clemens, K., Larsen, L., Zhang, M., Kuznetsov, Y., Bilanchone, V., Randall, A., Harned, A., DaSilva, R., Nagashima, K., McPherson, A., et al. (2011). The TY3 Gag3 Spacer Controls Intracellular Condensation and Uncoating. Journal of Virology 85, 3055–3066.

Dao, T.P., Martyniak, B., Canning, A.J., Lei, Y., Colicino, E.G., Cosgrove, M.S., Hehnly, H., and Castañeda, C.A. (2019). ALS-Linked Mutations Affect UBQLN2 Oligomerization and Phase Separation in a Position- and Amino Acid-Dependent Manner. Structure 27, 937–951.e5.

Deng, H.-X., Chen, W., Hong, S.-T., Boycott, K.M., Gorrie, G.H., Siddique, N., Yang, Y., Fecto, F., Shi, Y., Zhai, H., et al. (2011). Mutations in UBQLN2 cause dominant X-linked juvenile and adult-onset ALS and ALS/dementia. Nature 477, 211–215.

Dobin, A., Davis, C.A., Schlesinger, F., Drenkow, J., Zaleski, C., Jha, S., Batut, P., Chaisson, M., and Gingeras, T.R. (2013). STAR: ultrafast universal RNA-seq aligner. Bioinformatics 29, 15–21.

Ferraiuolo, L., Kirby, J., Grierson, A.J., Sendtner, M., and Shaw, P.J. (2011). Molecular pathways of motor neuron injury in amyotrophic lateral sclerosis. Nat Rev Neurol 7, 616–630.

Finley, D. (2009). Recognition and Processing of Ubiquitin-Protein Conjugates by the Proteasome. Annu. Rev. Biochem. 78, 477–513.

Frankish, A., Diekhans, M., Ferreira, A.-M., Johnson, R., Jungreis, I., Loveland, J., Mudge, J.M., Sisu, C., Wright, J., Armstrong, J., et al. (2019). GENCODE reference annotation for the human and mouse genomes. Nucleic Acids Research 47, D766– D773.

Gorrie, G.H., Fecto, F., Radzicki, D., Weiss, C., Shi, Y., Dong, H., Zhai, H., Fu, R., Liu, E., Li, S., et al. (2014). Dendritic spinopathy in transgenic mice expressing ALS/dementia-linked mutant UBQLN2. Proc Natl Acad Sci USA 111, 14524–14529.

Hjerpe, R., Bett, J.S., Keuss, M.J., Solovyova, A., McWilliams, T.G., Johnson, C., Sahu, I., Varghese, J., Wood, N., Wightman, M., et al. (2016). UBQLN2 Mediates Autophagy- Independent Protein Aggregate Clearance by the Proteasome. Cell 166, 935–949.

Hughes, C.S., Foehr, S., Garfield, D.A., Furlong, E.E., Steinmetz, L.M., and Krijgsveld, J. (2014). Ultrasensitive proteome analysis using paramagnetic bead technology. Mol Syst Biol 10, 757.

Itakura, E., Zavodszky, E., Shao, S., Wohlever, M.L., Keenan, R.J., and Hegde, R.S. (2016). Ubiquilins Chaperone and Triage Mitochondrial Membrane Proteins for Degradation. Mol Cell 63, 21–33.

Kelley, L.A., Mezulis, S., Yates, C.M., Wass, M.N., and Sternberg, M.J.E. (2015). The Phyre2 web portal for protein modeling, prediction and analysis. Nat Protoc 10, 845– 858.

Kim, S., Thaper, D., Bidnur, S., Toren, P., Akamatsu, S., Bishop, J.L., Colins, C., Vahid, S., and Zoubeidi, A. (2019). PEG10 is associated with treatment-induced neuroendocrine prostate cancer. Journal of Molecular Endocrinology 63, 39–49.

Kirchner, J., and Sandmeyer, S. (1993). Proteolytic processing of Ty3 proteins is required for transposition. J Virol 67, 19–28.

Klementieva, N.V., Lukyanov, K.A., Markina, N.M., Lukyanov, S.A., Zagaynova, E.V., and Mishin, A.S. (2016). Green-to-red primed conversion of Dendra2 using blue and red lasers. Chem. Commun. 52, 13144–13146.

Kvartsberg, H., Lashley, T., Murray, C.E., Brinkmalm, G., Cullen, N.C., Höglund, K., Zetterberg, H., Blennow, K., and Portelius, E. (2019). The intact postsynaptic protein neurogranin is reduced in brain tissue from patients with familial and sporadic Alzheimer’s disease. Acta Neuropathol 137, 89–102.

Larsen, L.S.Z., Beliakova-Bethell, N., Bilanchone, V., Zhang, M., Lamsa, A., DaSilva, R., Hatfield, G.W., Nagashima, K., and Sandmeyer, S. (2008). Ty3 Nucleocapsid Controls Localization of Particle Assembly. JVI 82, 2501–2514.

Le, N.T.T., Chang, L., Kovlyagina, I., Georgiou, P., Safren, N., Braunstein, K.E., Kvarta, M.D., Van Dyke, A.M., LeGates, T.A., Philips, T., et al. (2016). Motor neuron disease, TDP-43 pathology, and memory deficits in mice expressing ALS–FTD-linked UBQLN2 mutations. Proc Natl Acad Sci USA 113, E7580–E7589.

Lee, D.Y., and Brown, E.J. (2012). Ubiquilins in the crosstalk among proteolytic pathways. Biological Chemistry 393, 441–447.

Li, H., Handsaker, B., Wysoker, A., Fennell, T., Ruan, J., Homer, N., Marth, G., Abecasis, G., Durbin, R., and 1000 Genome Project Data Processing Subgroup (2009). The Sequence Alignment/Map format and SAMtools. Bioinformatics 25, 2078–2079.

Love, M.I., Huber, W., and Anders, S. (2014). Moderated estimation of fold change and dispersion for RNA-seq data with DESeq2. Genome Biol 15, 550.

Luo, W., and Brouwer, C. (2013). Pathview: an R/Bioconductor package for pathway- based data integration and visualization. Bioinformatics 29, 1830–1831.

Lux, H., Flammann, H., Hafner, M., and Lux, A. (2010). Genetic and Molecular Analyses of PEG10 Reveal New Aspects of Genomic Organization, Transcription and Translation. PLoS ONE 5, e8686.

Manktelow, E. (2005). Characterization of the frameshift signal of Edr, a mammalian example of programmed -1 ribosomal frameshifting. Nucleic Acids Research 33, 1553– 1563.

Marín, I. (2014). The ubiquilin gene family: evolutionary patterns and functional insights. BMC Evol Biol 14, 63.

Ono, R., Nakamura, K., Inoue, K., Naruse, M., Usami, T., Wakisaka-Saito, N., Hino, T., Suzuki-Migishima, R., Ogonuki, N., Miki, H., et al. (2006). Deletion of Peg10, an imprinted gene acquired from a retrotransposon, causes early embryonic lethality. Nat Genet 38, 101–106.

Pandya, N.J., Wang, C., Costa, V., Lopatta, P., Meier, S., Zampeta, F.I., Punt, A.M., Mientjes, E., Grossen, P., Distler, T., et al. (2021). Secreted retrovirus-like GAG- domain-containing protein PEG10 is regulated by UBE3A and is involved in Angelman syndrome pathophysiology. Cell Reports Medicine 2, 100360.

Pantano, L. (2021). DEGreport: Report of DEG analysis. R package version 1.30.0, http://lpatano.github.io/DEGreport/. (Bioconductor).

Pastuzyn, E.D., Day, C.E., Kearns, R.B., Kyrke-Smith, M., Taibi, A.V., McCormick, J., Yoder, N., Belnap, D.M., Erlendsson, S., Morado, D.R., et al. (2018). The Neuronal Gene Arc Encodes a Repurposed Retrotransposon Gag Protein that Mediates Intercellular RNA Transfer. Cell 172, 275–288.e18.

Peters, O.M., Ghasemi, M., and Brown, R.H. (2015). Emerging mechanisms of molecular pathology in ALS. J. Clin. Invest. 125, 1767–1779.

Riedl, J., Crevenna, A.H., Kessenbrock, K., Yu, J.H., Neukirchen, D., Bista, M., Bradke, F., Jenne, D., Holak, T.A., Werb, Z., et al. (2008). Lifeact: a versatile marker to visualize F-actin. Nat Methods 5, 605–607.

Saeki, Y. (2017). Ubiquitin recognition by the proteasome. J Biochem mvw091.

Sandmeyer, S.B., and Clemens, K.A. (2010). Function of a retrotransposon nucleocapsid protein. RNA Biology 7, 642–654.

Segel, M., Lash, B., Song, J., Ladha, A., Liu, C.C., Jin, X., Mekhedov, S.L., Macrae, R.K., Koonin, E.V., and Zhang, F. (2021). Mammalian retrovirus-like protein PEG10 packages its own mRNA and can be pseudotyped for mRNA delivery. Science 373, 882–889.

Sharkey, L.M., Safren, N., Pithadia, A.S., Gerson, J.E., Dulchavsky, M., Fischer, S., Patel, R., Lantis, G., Ashraf, N., Kim, J.H., et al. (2018). Mutant UBQLN2 promotes toxicity by modulating intrinsic self-assembly. Proc Natl Acad Sci USA 115, E10495– E10504.

Sharkey, L.M., Sandoval-Pistorius, S.S., Moore, S.J., Gerson, J.E., Komlo, R., Fischer, S., Negron-Rios, K.Y., Crowley, E.V., Padron, F., Patel, R., et al. (2020). Modeling UBQLN2-mediated neurodegenerative disease in mice: Shared and divergent properties of wild type and mutant UBQLN2 in phase separation, subcellular localization, altered proteostasis pathways, and selective cytotoxicity. Neurobiology of Disease 143, 105016.

Steplewski, A., Krynska, B., Tretiakova, A., Haas, S., Khalili, K., and Amini, S. (1998). MyEF-3, a Developmentally Controlled Brain-Derived Nuclear Protein Which Specifically Interacts with Myelin Basic Protein Proximal Regulatory Sequences. Biochemical and Biophysical Research Communications 243, 295–301.

Suzuki, R., and Kawahara, H. (2016). UBQLN4 recognizes mislocalized transmembrane domain proteins and targets these to proteasomal degradation. EMBO Rep 17, 842– 857.

Tsubaki, H., Tooyama, I., and Walker, D.G. (2020). Thioredoxin-Interacting Protein (TXNIP) with Focus on Brain and Neurodegenerative Diseases. IJMS 21, 9357.

Vaquero-Garcia, J., Barrera, A., Gazzara, M.R., González-Vallinas, J., Lahens, N.F., Hogenesch, J.B., Lynch, K.W., and Barash, Y. (2016). A new view of transcriptome complexity and regulation through the lens of local splicing variations. ELife 5, e11752.

Vaquero-Garcia, J., Aicher, J.K., Jewell, P., Gazzara, M.R., Radens, C.M., Jha, A., Green, C.J., Norton, S.S., Lahens, N.F., Grant, G.R., et al. (2021). RNA splicing analysis using heterogeneous and large RNA-seq datasets (Bioinformatics).

Vijayakumar, U.G., Milla, V., Cynthia Stafford, M.Y., Bjourson, A.J., Duddy, W., and Duguez, S.M.-R. (2019). A Systematic Review of Suggested Molecular Strata, Biomarkers and Their Tissue Sources in ALS. Front. Neurol. 10, 400.

Volff, J.-N. (2006). Turning junk into gold: domestication of transposable elements and the creation of new genes in eukaryotes. Bioessays 28, 913–922.

Whiteley, A.M., Prado, M.A., Peng, I., Abbas, A.R., Haley, B., Paulo, J.A., Reichelt, M., Katakam, A., Sagolla, M., Modrusan, Z., et al. (2017). Ubiquilin1 promotes antigen- receptor mediated proliferation by eliminating mislocalized mitochondrial proteins. Elife 6.

Whiteley, A.M., Prado, M.A., de Poot, S.A.H., Paulo, J.A., Ashton, M., Dominguez, S., Weber, M., Ngu, H., Szpyt, J., Jedrychowski, M.P., et al. (2021). Global proteomics of Ubqln2-based murine models of ALS. J. Biol. Chem. 296, 100153.

Williams, K.L., Warraich, S.T., Yang, S., Solski, J.A., Fernando, R., Rouleau, G.A., Nicholson, G.A., and Blair, I.P. (2012). UBQLN2/ubiquilin 2 mutation and pathology in familial amyotrophic lateral sclerosis. Neurobiology of Aging 33, 2527.e3–2527.e10.

Yu, G., Wang, L.-G., Han, Y., and He, Q.-Y. (2012). clusterProfiler: an R Package for Comparing Biological Themes Among Gene Clusters. OMICS: A Journal of Integrative Biology 16, 284–287.

Yu, G., Wang, L.-G., Yan, G.-R., and He, Q.-Y. (2015). DOSE: an R/Bioconductor package for disease ontology semantic and enrichment analysis. Bioinformatics 31, 608–609.

Zhang, D., Raasi, S., and Fushman, D. (2008). Affinity Makes the Difference: Nonselective Interaction of the UBA Domain of Ubiquilin-1 with Monomeric Ubiquitin and Polyubiquitin Chains. Journal of Molecular Biology 377, 162–180.

Zheng, T., Yang, Y., and Castañeda, C.A. (2020). Structure, dynamics and functions of UBQLNs: at the crossroads of protein quality control machinery. Biochemical Journal 477, 3471–3497.

