## Supplemental Figures for "UBQLN2 restrains the domesticated retrotransposon PEG10 to maintain neuronal health in ALS"

**Supplementary Table 1: RNA-Seq results and analysis.** Differentially expressed genes (DEGs) in a pairwise DESeq2 analysis with control plasmid are shown in unique tabs (A-C) for different PEG10 constructs. Overrepresentation analysis of pathways of genes significantly upregulated with a lfc cutoff of 0.5 and p value of 0.05 is shown for each unique PEG10 construct (D-F). Finally, splice alterations (differentially spliced genes, DSGs) using MAJIQ analysis are summarized with a deltapsi threshold of 0.1 and p value 0.05 are shown in (G-I).

**Supplementary Table 2: Global proteomics of human spinal cord.** Human sample metadata is summarized in (A). Spectrometer parameters are summarized in (B). Organized global proteomic results are shown in (C), with non-neurological controls (NNC) in blue, sALS patients in yellow, and UBQLN2 fALS in orange. Log2 ALS/NNC and inverse log T-test are shown in green. Reverse peptides, potential contaminants, and proteins identified by only one peptide were excluded for quantification analysis in Figure 4. Peptide-level data for PEG10 is shown in (D). Gag-derived peptides are shown in pink, and pol-derived peptides are shown in blue.

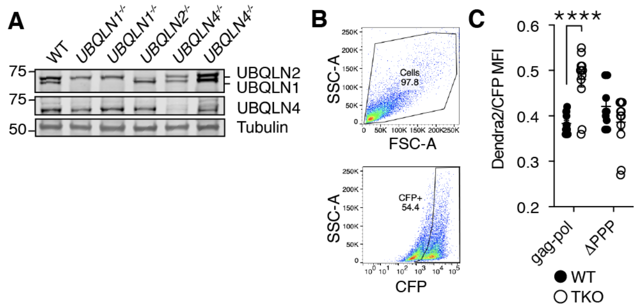


**Supplemental Figure S1: PEG10 and UBQLN2 specificity screening strategy. (A)** Human ESCs had individual *UBQLN* genes deleted by CRISPR gene editing and clones were probed by western blot for each individual UBQLN. n = 3 representative blots. Duplicate cell lines represent parallel clones generated from CRISPR gene editing process. **(B)** Example gating strategy of abundance reporter test. 'Cells' are gated (left), then gated on CFP+ population (middle, with Dendra-green fluorescence of CFP+ population shown at right). A custom parameter of Dendra/CFP is then generated on a per-cell basis and the geometric mean fluorescence intensity (MFI) of this custom parameter is averaged per sample. **(C)** Dendra-green over CFP MFI ratios for PEG10 (data from Figure 1) and ΔPPR PEG10 in WT (black circles) and TKO (outline circles) cells. Shown is mean ± SEM from four independent experiments with triplicate transfection wells. Unpaired comparison between WT and TKO Dendra2/CFP ratios was performed by Student's t-test.

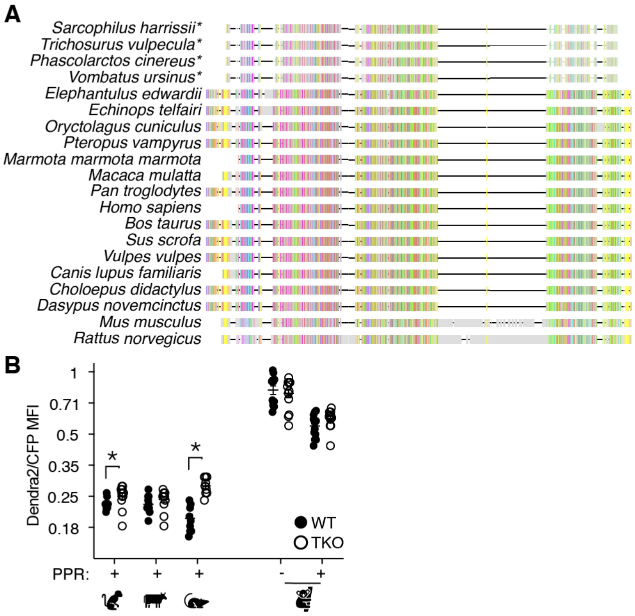


**Supplemental Figure S2: Evolutionary analysis of PEG10-UBQLN2 relationship. (A)** Amino acid alignment of mammalian PEG10 from a diversity of mammalian species showing general conservation and lack of C-terminal polyproline domain in marsupials (starred). Colors represent conservation of aligned amino acids; proline is yellow. **(B)** Dendra-green over CFP MFI ratios for PEG10 from various mammalian species in WT (filled circles) and TKO (open circles) cells. Shown is mean ± SEM from four independent experiments, with triplicate transfection wells for each. Unpaired comparison between WT and TKO Dendra2/CFP ratios was performed by Student's t-test.

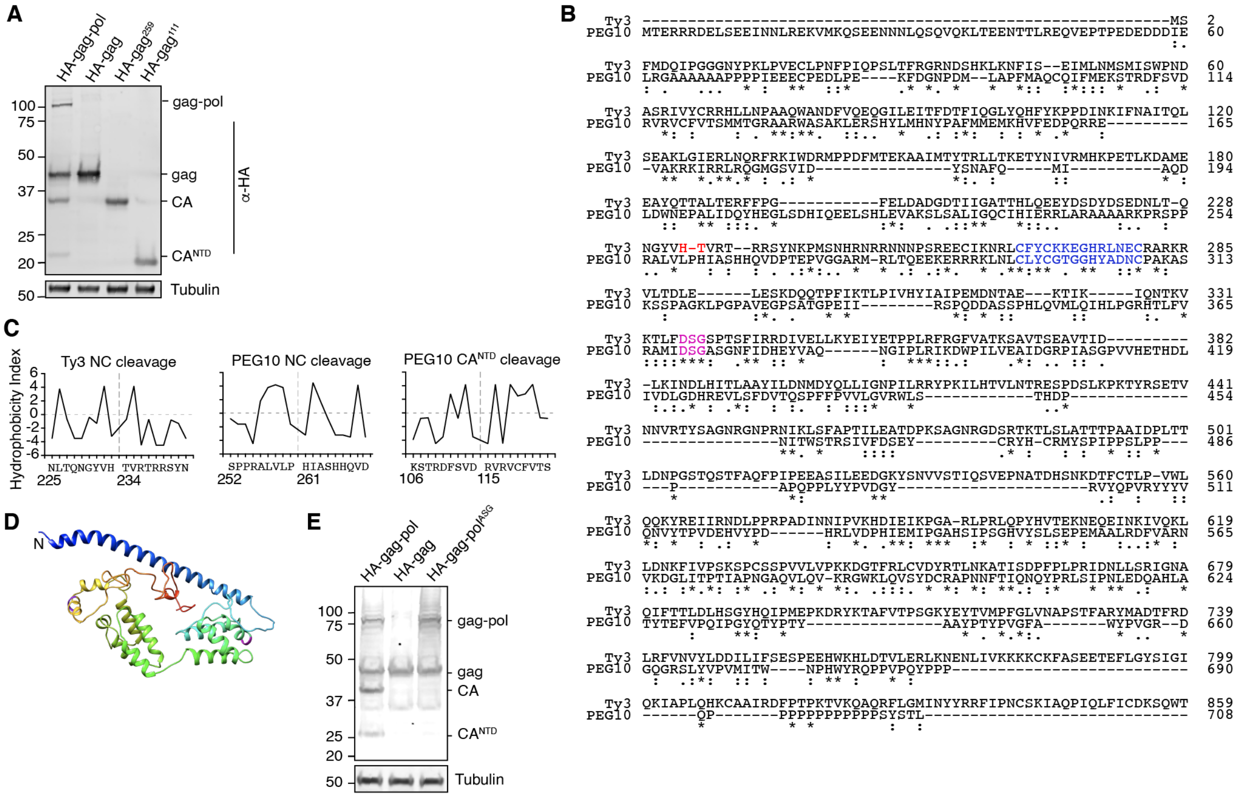


**Supplemental Figure S3: PEG10 self-cleavage locations and comparison to Ty3.** **(A)** HA-tagged, C-terminally truncated forms of PEG10 were expressed in cells to approximate self-cleavage sites via molecular weight. n = 3 independent experiments. **(B)** Alignment of human PEG10 with the ancestral Ty3 retrotransposon from *Saccharomyces cerevisiae*. Conserved active site of protease is in magenta; CCHC domain is in blue, and a known Ty3 self-cleavage site is in red. Homology is indicated underneath alignment according to ClustalOmega parameters (Sievers et al., 2011). **(C)** Kyte-Doolittle (Kyte and Doolittle, 1982) (K-D) hydropathy analysis of Ty3 (left, similar to that in Kirchner and Sandmeyer (Kirchner and Sandmeyer, 1993)) and PEG10 nucleocapsid (right) and $CA^{NTD}$ (bottom) cleavages. Hydropathy measurements for each amino acid were plotted on a scale from P9 to P9'. Amino acid locations are shown below the amino acid alignment. Estimated cleavage locations by molecular weight in Figure 3D are close in proximity to those estimated by K-D analysis. **(D)** Structure prediction of PEG10 gag (blue N-terminus) shows that estimated cleavage sites appear accessible by protease. AA114-115 (purple) and AA260-261 (magenta) are highlighted to show estimated cleavages. **(E)** Western blot of accompanying cell lysate from VLP preparations. In parallel to media harvest for VLP isolation, cell lysate was prepared to probe relative quantities of PEG10 production and presence of cleavage products. n = 3 independent experiments.

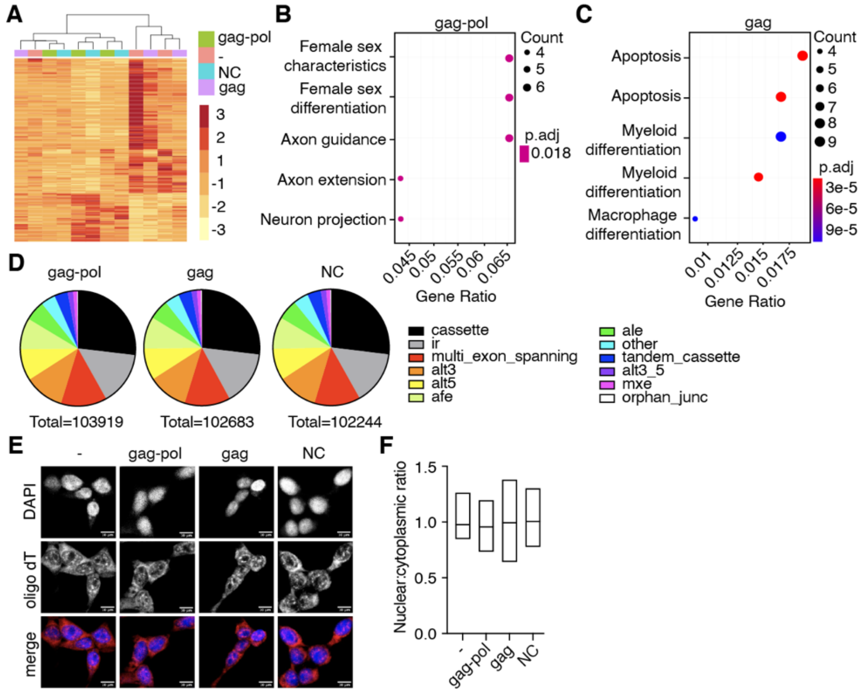


**Supplemental Figure S4: Transcriptional effects of PEG10 nucleocapsid transfection resemble gag-pol, but not gag. (A)** Heatmap showing Euclidean clustering of transfection constructs with top 200 altered genes between gag-pol and negative control plasmid. **(B-C)** Top gene expression changes by GO-term enrichment analysis. The top five GO-terms enriched upon comparison of gag-pol (B) and gag (C) to control cells are shown with enriched GO-term on the left. Adjusted p value is shown by color, and size of datapoint reflects the number of genes enriched in the pathway. **(D)** Splice pattern changes upon PEG10 overexpression compared to control. Nucleocapsid expression does not alter patterns of splice alteration compared to gag-pol or gag expression. Splice changes were quantified and classified by MAJIQ analysis (Vaquero-Garcia et al., 2016). Total splice alteration counts are shown below pie charts. ir = intron retention; afe/ale = alternative first/last exon; mxe = mutually exclusive exons. **(E)** Oligo-dT FISH showing no changes to bulk mRNA trafficking upon PEG10 overexpression. Scale bar 10 μm. Shown are representative images from 10 recorded fields of view. n=3 independent experiments. **(F)** Quantification of oligo-dT signal in nucleus versus cytosol for each condition imaged in (E) showing no changes to mRNA trafficking upon PEG10 overexpression. Quantification was performed on a minimum of 60 images from each condition.

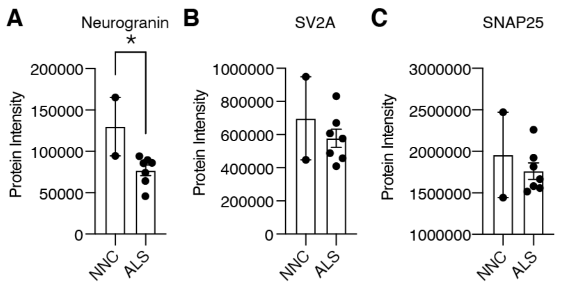


**Supplemental Figure S5: Putative biomarkers of neurodegeneration in ALS patient tissue. (A-C)** Abundance of neurogranin (A), SV2A (B), and SNAP25 (C), in human spinal cord. Mean ± SEM is shown and plotted for each marker. A significant decrease in neurogranin was detected between non-neurological controls NNC (n = 2) and all ALS patients (n = 7) by unpaired t-test.

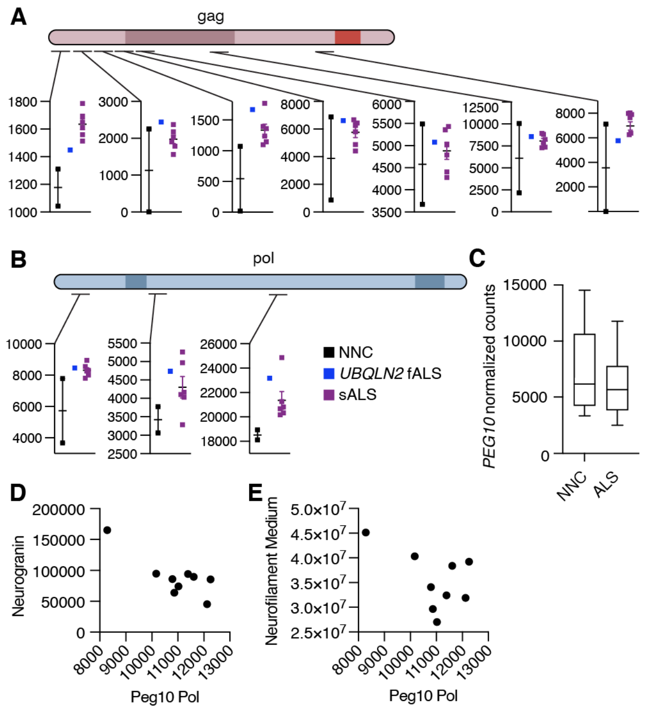


**Supplemental Figure S6: Abundance of PEG10 is changed in ALS at the protein, but not the mRNA, level. (A-B)** Schematic of PEG10 protein from gag (A) and pol (B) with regions highlighted where peptides were quantified by global proteomic analysis. More detail on quantified peptides can be found in Supplementary Data Table 2. Black: Non-neurological control, blue: *UBQLN2*-fALS, purple: sporadic ALS. Peptide abundance as measured by mass spectrometer is shown on *y* axis, with mean ± SEM of replicates. **(C)** RNA-Seq counts of *PEG10* from post-mortem lumbar spinal cord tissue of patients with Classical ALS or Non-Neurological Controls (NNC). **(D)** Quantification of neurogranin (*y* axis) and PEG10 pol peptides (*x* axis) are plotted for all 9 spinal cord samples to demonstrate the relationship between the two markers. **(E)** Quantification of neurofilament medium (*y* axis) and PEG10 pol peptides (*x* axis) are plotted for all 9 spinal cord samples to demonstrate the relationship between the two markers.
